# Progesterone specifically dampens disease-associated T_H_1- and T_H_17-related immune responses during T cell activation *in vitro*

**DOI:** 10.1101/2020.11.05.370700

**Authors:** Sandra Hellberg, Johanna Raffetseder, Olof Rundquist, Rasmus Magnusson, Georgia Papapavlou, Maria C. Jenmalm, Jan Ernerudh, Mika Gustafsson

## Abstract

The changes in progesterone (P4) levels during and after pregnancy coincide with the temporary improvement and worsening of several autoimmune diseases like multiple sclerosis (MS) and rheumatoid arthritis (RA). Most likely immune-endocrine interactions play a major role in these pregnancy-induce effects. In this study, we used next generation sequencing to investigate the direct effects of P4 on CD4^+^ T cell activation, of central importance in pregnancy and disease. We found that P4 had a profound dampening effect on T cell activation, altering the gene and protein expression profile and opposing many of the changes induced during the activation. The transcriptomic changes induced by P4 were significantly enriched for genes associated with diseases known to be modulated during pregnancy such as MS, RA and psoriasis. The T_H_1-and T_H_17-associated transcription factors STAT1 and STAT3 were significantly downregulated by P4 and their downstream targets were significantly enriched among the diseases-associated genes. Several of these genes included well-known and disease-relevant cytokines, such as IL-12β, CXCL10 and OSM, that were further validated also at the protein level using proximity extension assay. Our results extend the previous knowledge of P4 as an immune regulatory hormone and supports its importance during pregnancy for regulating potentially detrimental immune responses towards the semi-allogenic fetus. Further, our results also point toward a potential role for P4 in the pregnancy-induced disease immunomodulation, suggestively through dampening of T_H_1 and T_H_17-associated immune responses and highlights the need for further studies evaluating P4 as a future treatment option.

## INTRODUCTION

Pregnancy represents a unique immunological condition as the maternal immune system is able to tolerate the presence of the semi-allogenic fetus. This immunological tolerance is thought to arise from extensive immune and endocrine alterations induced during pregnancy^1, 2^ orchestrated by the increased levels of the steroid hormone progesterone (P4), which is essential for the establishment and maintenance of pregnancy.^3–5^ Accordingly, low levels of P4 have been associated with several pregnancy complications^6–8^ and, in fact, treatment with the P4-receptor antagonist Mifepristone (RU486) results in cessation of pregnancy,^9^ further supporting the importance of P4 in pregnancy. Interestingly, therapeutic use of P4 in pregnancy has been shown to reduce the risk of preterm birth in certain risk groups,^10^ highlighting a potential use of P4 as treatment in the pregnancy complications.

The role of P4 has been mainly related to its influence on myometrial homeostasis and remodeling.^11^ However, an important immune-modulatory role of P4 *in vivo* is indicated by the pregnancy-related improvement and subsequent worsening of inflammatory diseases, such as multiple sclerosis (MS) and rheumatoid arthritis (RA), which coincides with the time points during and after pregnancy when P4 levels are the highest and lowest, respectively.^12–14^ In addition, differences in immune responses related to the high and low P4 levels during the menstrual cycle have also been observed.^15, 16^ Previous studies regarding immune-endocrine interactions have to a large extent focused on how estrogen influences the immune system. Interestingly, estrogen levels increase and decrease during pregnancy in a similar way as P4. However, estrogen seems to have both immune regulatory and immune-activating properties.^17^ Indeed, estrogen has been suggested as a major factor explaining the increased occurrence of autoimmune disease in women.^18^ On the other hand, *in vitro* effects of P4 on different immune cell populations support a pivotal role for P4 in regulating immune responses that could be central for promoting fetal tolerance.^19–24^ Furthermore, a role for P4 as a potent immunosuppressor is supported by *in vivo* studies showing involvement of P4 in response to allogenic and xenogeneic transplantation ‘ and in graft rejections in humans.

CD4^+^ T cells are central in the immune system, serving as chief regulators of immunity and tolerance.^29^ The importance of CD4^+^ T cells during pregnancy is evident by their relative exclusion at the fetal-maternal interface in order to limit potentially detrimental activation,^30^ whereas regulatory T cells are enriched, thereby further limiting harmful T cell responses.^31, 32^ In parallel, limiting of CD4^+^ T cell activation is an essential aspect in T cell-mediated diseases as aberrant activation of autoreactive CD4^+^ T cells is a central mechanism in the disease pathogenesis.^33^ Thus, regulation of CD4^+^T cells is a common denominator that could both prevent unwanted maternal immune responses against the fetus, and also explain improvement of inflammatory diseases during pregnancy. Interestingly, P4 has been shown to limit CD4^+^ T cell activation.^34–36^ However, in-depth analysis of the precise effects of P4 on human CD4^+^T cell activation and its potential involvement in disease modulation during pregnancy is still lacking.

We here report in-depth RNA sequencing data demonstrating a profound direct effect of P4 on T cell activation. More specifically, P4 induced large transcriptomic changes, most prominently downregulatory, on immune-associated genes and pathways and interestingly, these genes were significantly enriched for genes associated with diseases known to be modulated during pregnancy including MS and RA. These changes were mainly related to STAT1 and STAT3 and their downstream targets, suggesting a possible involvement of P4 in dampening T_H_1- and T_H_17-associated immune responses of relevance for disease. These findings suggest that P4 could be involved in mediating the pregnancy-induced improvement of certain inflammatory diseases and constitute a potential avenue for future treatment options.

## RESULTS

### Progesterone dampens T cell activation and affects the transcriptomic profile in activated CD4^+^ T cells

In order to examine the effect of P4 on CD4^+^ T cells and T cell activation, we established an *in vitro* model where primary human CD4^+^ T cells were first pre-incubated with P4 prior to activation, reflecting the *in vivo* situation where the T cells would constantly be exposed to P4 prior to antigen challenge. The cells were subsequently cultured unactivated or with the commonly used combined activation through the T cell receptor (anti-CD3) and co-stimulatory CD28 (anti-CD28) in the presence or absence of P4 for 6, 24 and 72 hrs (**Fig. 1a**). These time points were chosen in order to mainly capture the influence of P4 on the cellular and transcriptomic events involved in T cell activation leading up to clonal expansion and differentiation. We used a low-to-moderate level of stimulation, as measured by the proportion of activated T cells expressing surface activation markers (6 hrs CD69, mean± SD unactivated: 0.2%+0.04, activated: 15%± 9.5 and 24 hrs CD69, unactivated: 0.2%+0.07, activated: 20%+9.4; CD25, unactivated: 6.5%±1.4, activated: 16%±4.7) (**Fig. 1b-d**). Activation of CD4^+^T cells in the presence of different concentrations of P4 decreased the level of activation of the cells in a dose-dependent manner, with reduced proportion of both CD69 and CD25 expressing cells, where exposure to the highest concentration of P4 (50 μM) resulted in the largest overall decrease (6 hrs: CD69^+^ *p*<0.010, 0.4±0.1 fold change compared to activation without P4 *p*<0.0001; 24 hrs: CD69^+^ *p*<0.001, 0.4±0.1 fold change *p*<0.0001 and CD25^+^ *p*<0.0001, 0.5+0.1 fold change *p*<0.0001) **(Fig. 1b-d and Supplementary Fig. 1)**. Next, we performed transcriptomic analysis using RNA sequencing (RNA-Seq) on CD4^+^ T cells activated for 6 and 24 hrs in the presence (50 μM P4) or absence of P4 along with unactivated cells, to obtain an in-depth picture of how T cell activation is affected by P4. To get an initial global understanding of the transcriptomic changes, we performed unsupervised clustering of the samples using multidimensional scaling, which showed clear differences between the cells activated in the presence or absence of P4, as well as in comparison to unactivated cells **(Fig. 1e-f)**. Indeed, differential expression analysis revealed that P4 induced significant changes in the gene expression profile of the activated CD4^+^ T cells at both 6 and 24 hrs (as compared to activation in the absence of P4), hereon referred to as P4 response genes. In total, P4 exposure resulted in 4276 differentially expressed genes (DEGs, false discover rate (FDR) ≤0.05; 2080 up-regulated and 2196 down-regulated) at 6 hrs and 4756 DEGs (2340 up-regulated and 2416 down-regulated) at 24 hrs **(Fig. 1g-h)**.

**Fig. 1.**
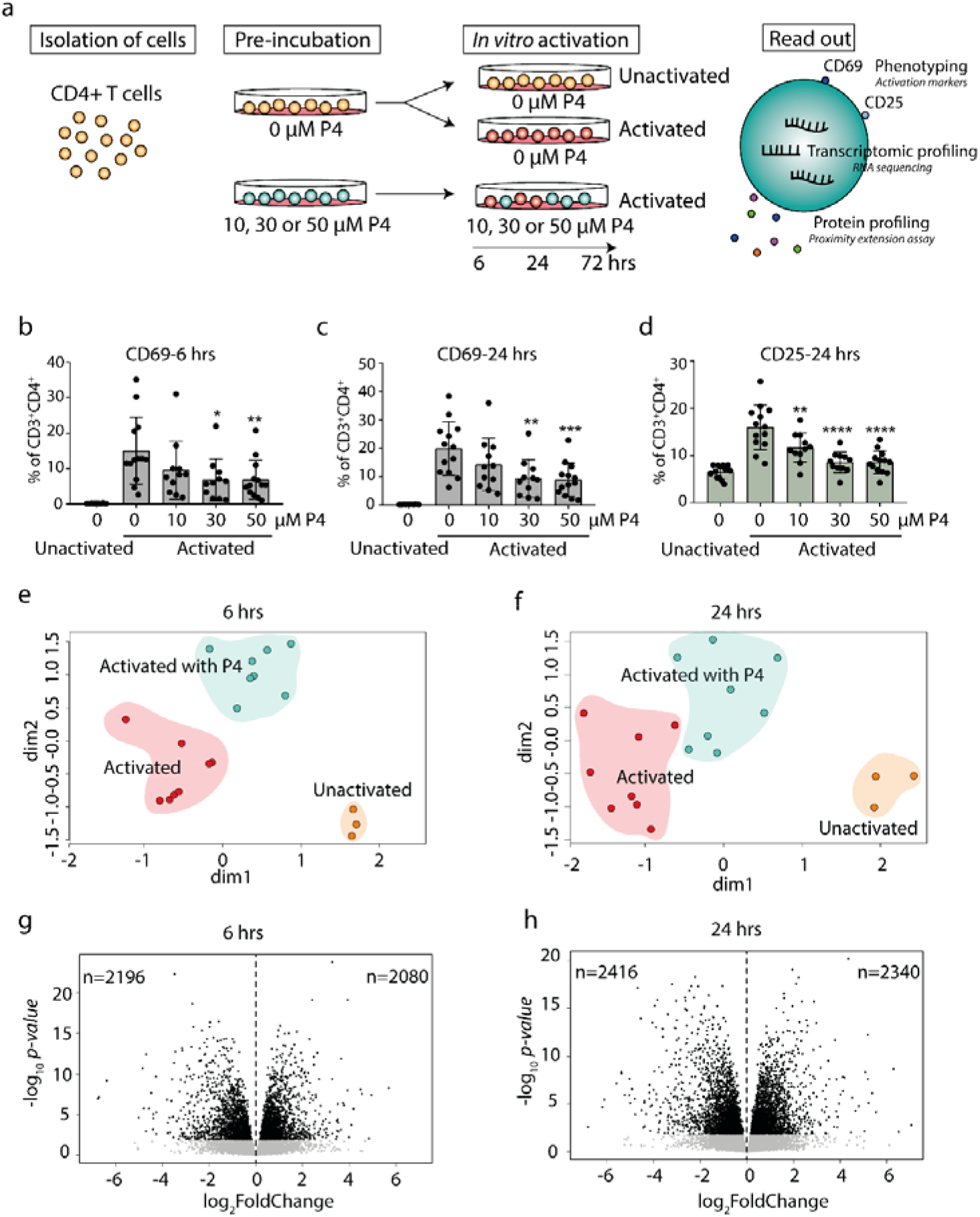
Progesterone dampens T cell activation and induces transcriptomic changes in activated CD4^+^ T cells. **a** Primary human CD4^+^ T cells were isolated from healthy non-pregnant women and pre-incubated with or without different concentrations (10, 30 and 50 μM) of P4 for 20 hrs and then cultured unactivated or activated *in vitro* of plate-bound anti-CD3 and anti-CD28 antibodies in the presence or absence of P4 for 6-24-72 hrs. Samples cultured for 72 hrs were only used for measurement of secreted proteins. The effect of P4 on T cell activation was evaluated by flow cytometry, RNA sequencing and proximity extension assay of secreted proteins in culture supernatants, b The effect of P4 on T cell activation markers CD69 (6 and 24 hrs) and CD25 (24 hrs only) was analysed by flow cytometry (n=11-13). Bar graphs shows the percentages of CD4^+^ T cells expressing the T cell activation markers. Mean ± standard deviations are shown. *p≤0.05, **p≤0.01, ***p≤0.001, ****p≤0.0001. **c-d** Multidimensional scaling analysis of gene expression data generated by RNA-sequencing (n=3 unactivated, n=8 for activated with and without 50 μM P4). The groups were highlighted by background color for schematic purposes only. **e-f** Volcano plots of the transcriptomic analysis of the differentially expressed genes in CD4^+^ T cells activated in the presence of P4 as compared to absence of P4. Black dots FDR≤0.05. P4, progesterone.

### Gene set enrichment analysis shows a predominantly downregulatory effect of progesterone on immune-related pathways affecting particularly genes associated with T cell activation

To gain further insight into the functional significance of the P4 response genes, we performed a gene set enrichment analysis (GSEA). Several immune-related pathways associated with and downstream of T cell signaling were significantly downregulated by P4 at both 6 and 24 hrs, for example T cell receptor signaling, JAK-STAT signaling, cytokine-cytokine receptor interactions, as well as many immune-associated disease pathways **(Fig. 2a-b and Supplementary Table 1).** At 24 hrs, several pathways related to T cell differentiation (Th1, Th2 and Th17 differentiation) were also affected by P4. Strikingly, GSEA showed very few pathways to be upregulated by P4 even though there was an almost equal number of P4 response genes that were up- and downregulated. Furthermore, none of the upregulated pathways were immune-related **(Supplementary Table 1).**

**Fig. 2.**
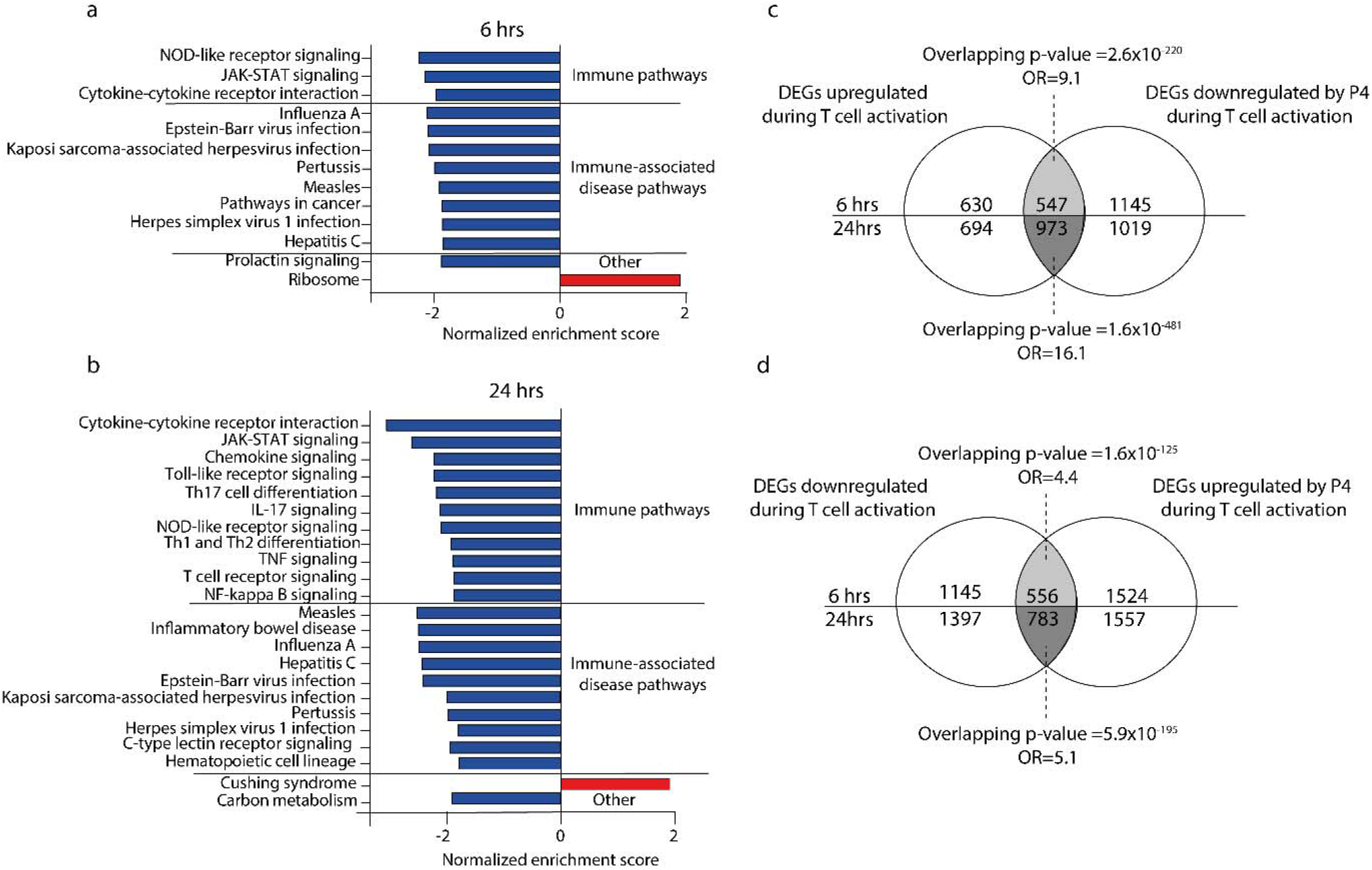
Many immune-related pathways are down-regulated in CD4^+^ T cells activated in the presence of progesterone. **a-b** Bar graphs showing the normalized enrichments score on the x-axis from the KEGG gene set enrichment analysis of the P4 response genes (DEGs comparing activation in the presence of P4 as compared to activation alone) at 6 and 24 hrs. All pathways have an FDR adjusted p-value <0.05. **c-d** Venn diagrams of the overlapping DEGs between up- and down-regulated genes in T cells activated in the presence or absence of 50 μM P4. DEGs, differentially expressed genes: OR, odds ratio: P4, progesterone.

Since P4 was found by us and by others^24, 35^ to have a dampening effect on T cell activation, we investigated if the changes induced by P4 were related to genes involved in the T cell activation process itself. We therefore assessed if P4 could counteract the changes induced during activation by analyzing the overlap between the P4 response genes and genes involved in the T cell activation, *i.e*. genes being differentially expressed in activated *versus* unactivated T cells. Indeed, there was a significant overlap between the DEGs upregulated during T cell activation and genes downregulated by P4 (6 hrs: odds ratio (OR): 9.1, p=2.6×10^-220^; 24 hrs: OR:16.1, p=1.6×10^-481^; **Fig. 2c**) and between the DEGs downregulated during T cell activation and upregulated by P4 (6 hrs: OR: 4.4, p=1.6×10^-125^; 24 hrs: OR: 5.1, p=5.9×10^-195^; **Fig. 2d**), demonstrating that P4 significantly affects genes related to the actual T cell activation by opposing some of the changes induced during activation. Conversely, very few genes showed the same directionality comparing P4 and T cell activation alone (data not shown).

### The dampening effect of P4 on gene expression of CD4^+^ T cells is evident also at the secreted protein level

To investigate if P4 also had a effect at the proteomics level, we performed a screening of 92 inflammation-related proteins in culture supernatants collected at 6, 24 and 72 hrs by using a proximity extension assay (PEA) with high sensitivity and specificity.^37^ At 6 hrs, only 24 out of the 92 proteins were detectable, of which 14 proteins were differentially expressed between CD4+ T cells cultured in the presence *versus* in the absence of P4 (**Fig. 3a** and **Supplementary Table 2**).

**Fig. 3.**
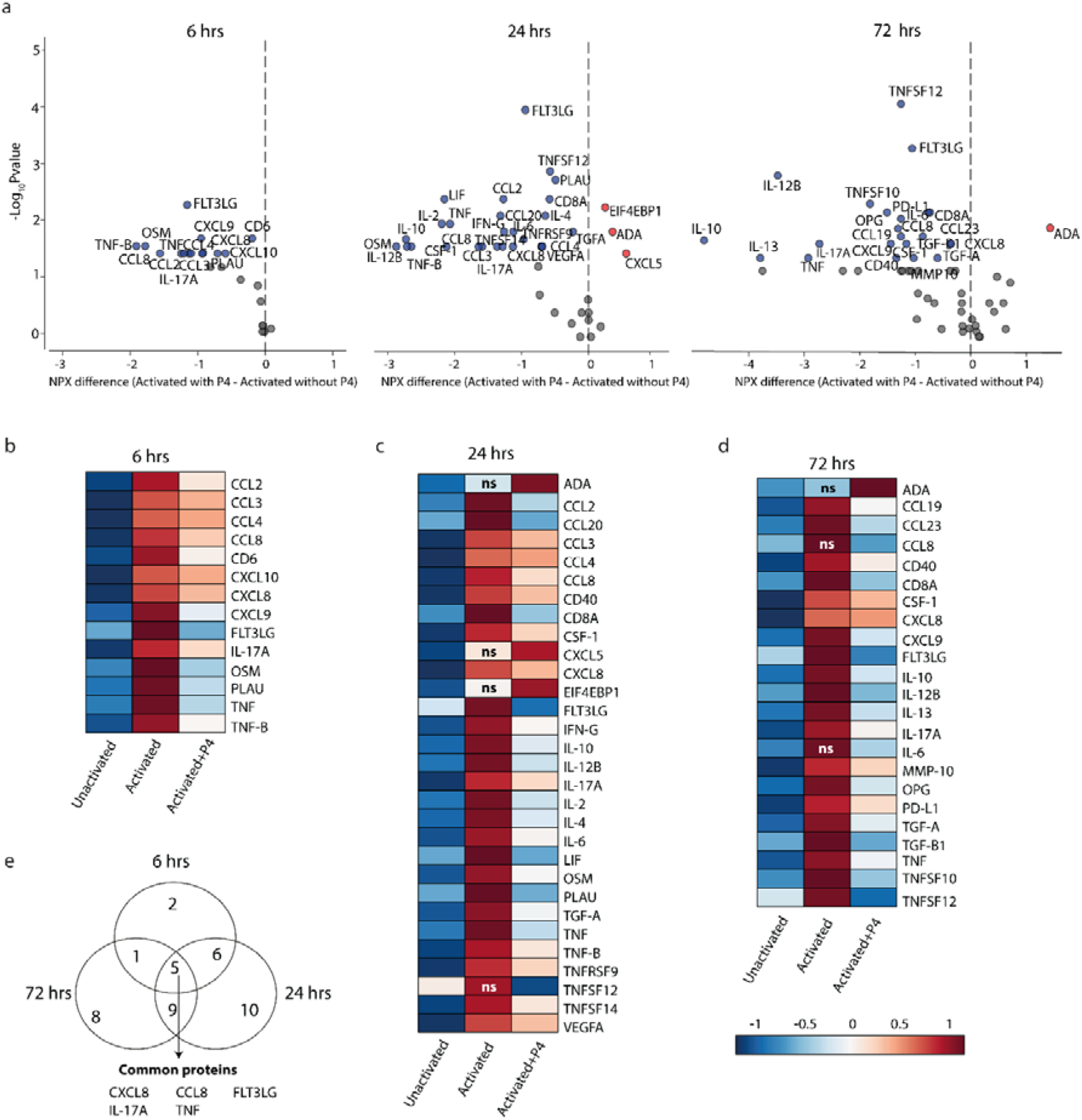
P4 significantly downregulates protein expression in culture supernatants. A panel of inflammation-related proteins were measured in supernatants collected from CD4^+^ T cells that were cultured unactivated or activated with or without 50 μM of P4. **a.** Volcano plots of differentially expressed proteins (DEPs) at 6, 24 and 72 hrs comparing activation in the presence or absence of P4. DEPs were determined using Friedman test and Benjamini-Hochberg. Data is presented as difference in the normalized protein expression values (NPX) provided by the manufacturer. Blue dots DEPs downregulated by P4, red dots upregulated DEPs and grey dots FDR≤0.05. **b-d.** Heat maps of the differentially expressed proteins comparing activation in the presence of P4 as compared to activation alone. ns are proteins that were not significantly different between activation alone as compared to unactivated. Median NPX values are shown in the figure. ns= non-significant. d. Venn diagram displaying the number of unique and common DEPs between the different time points.

At 24 hrs, 39 out of the 92 proteins were detectable and 28 (72%) of those were differentially expressed and at 72 hrs, 65 proteins were detectable but only 23 (35%) were differentially expressed (**Fig. 3a** and **Supplementary Table 2**). Consistent with the transcriptomic findings, most differentially expressed proteins (DEPs) were significantly lower in supernatants collected from T cells that had been activated in the presence of P4 as compared to activation alone. Only three proteins (ADA, CXCL5 and EIF4EBP1) were upregulated at any one time point. Also at the protein level, P4 seemingly opposed the changes induced during activation, where most proteins that were significantly upregulated during T cell activation (as compared to unactivated cells) were downregulated by P4 (**Fig. 3b-d**), except ADA (24 and 72 hrs), CCL8 (72hrs), CXCL5 (24 hrs), EIF4EBP1 (24 hrs), IL-6 (72 hrs) and TNFSF12 (24 hrs). There were five proteins that were consistently downregulated at all three time points: CXCL8, CCL8, IL-17A, TNF and FLT3LG (**Fig. 3d**). The dampening effect of P4 on immune-related processes at the transcriptomic levels is thus further supported by its apparent downregulatory effect on inflammation-related proteins.

### Progesterone significantly downregulates genes associated with immune-mediated diseases that are modulated during pregnancy

The fact that P4 levels during pregnancy coincide with the clinical changes observed in several immune-mediated diseases, prompted us to investigate if the genes affected by P4, *i.e*. the P4 response genes, were related to known disease genes. To this end, we used disease-associated genes derived from DisGeNET^38^ for seven diseases that have previously been shown to be altered during pregnancy^39^: Hashimoto’s disease, Graves’ disease, MS, psoriasis, RA, systemic lupus erythematosus(SLE) and systemic sclerosis. We combined the P4 response genes at both 6 and 24 hrs to capture if P4 affected disease-relevant genes, resulting in a total of 2563 downregulated and 2403 upregulated genes that were differentially expressed with P4 over the course of 24 hrs. In line with the previous finding of a much more pronounced downregulatory effect of P4 on immune-related genes, the downregulated P4 response genes were significantly enriched for genes associated with all seven diseases; Hashimoto’s disease (n=51, OR:2.6, p=1.6×10^-7^), Graves’ disease (n=73, OR: 2.0, p=2.3×10^-6^), psoriasis (n=134, OR: 1.4, p=5.8×10^-4^), MS (n=201, OR: 1.6, p=7.0×10^-8^), RA (n=270, OR:1.3, p=6.8×10^-4^), SLE (n=212, OR: 1.7, p=1.4×10^-9^) and systemic sclerosis (n=96, OR:1.5, p=8.5×10^-4^) **(Fig. 4a, Supplementary Table 3)** whereas the upregulated P4 response genes showed no significant enrichment for any disease-associated genes **(Fig. 4a)** Thus, only the downregulated P4 response genes were considered for the subsequent analysis. There was a total of 17 shared genes between all seven diseases that were significantly downregulated by P4: *BCL2, CXCL10, FAS, FOXP3, ICAM1, IL10, IL17A, IL1A, IL1B, IL21, IL22, IL23R, IL2R, ISG20, RBM45, TGFB1* and *TNF* **(Fig. 4a, Supplementary Table 3)**, highlighting potential central genes in the disease pathogenesis.

**Fig. 4.**
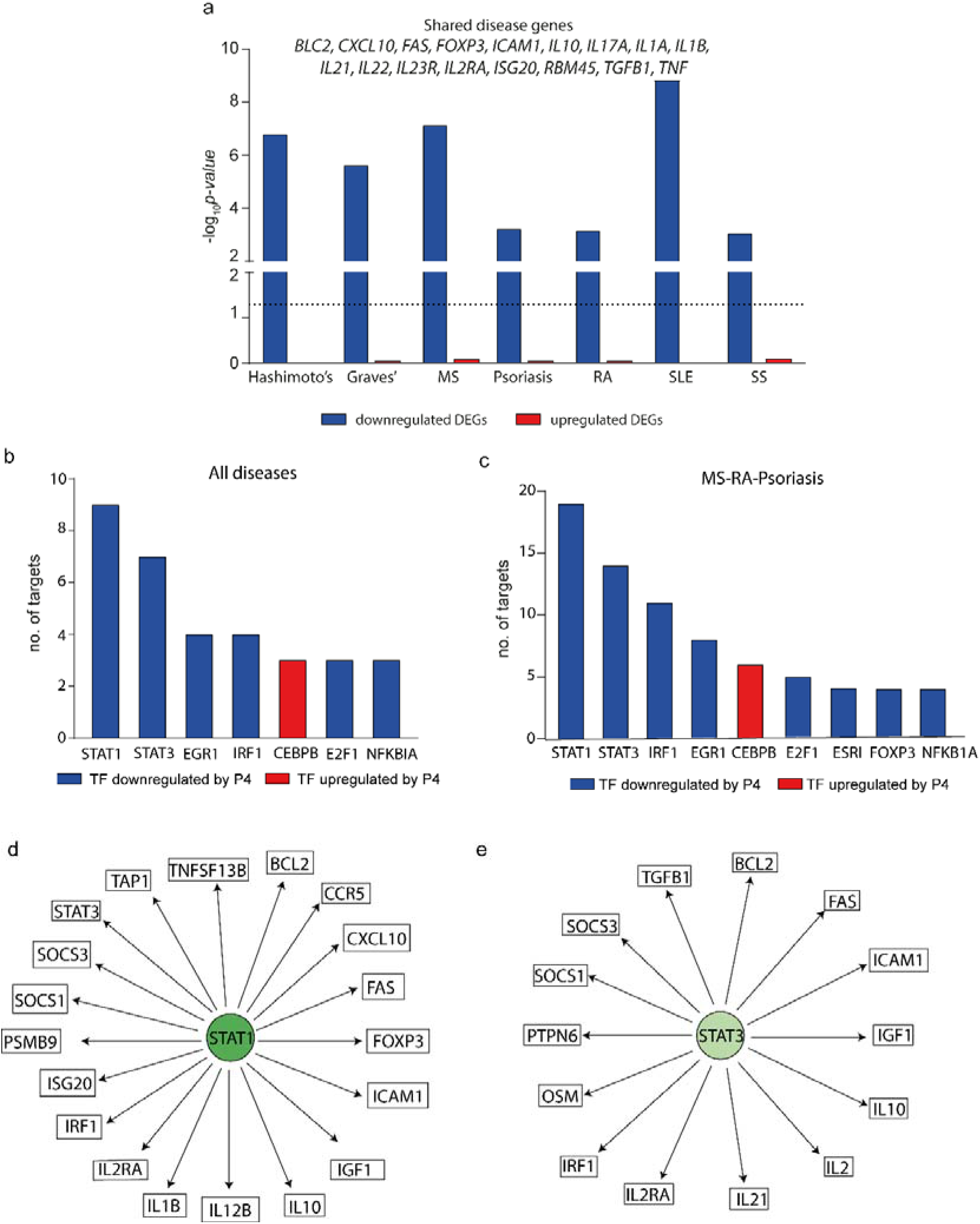
Progesterone downregulates disease-associated genes for several autoimmune diseases particularly through *STAT1* and *STAT3*. The effect of P4 on disease-associated genes and upstream transcription factors was interrogated using known disease genes (derived from DisGeNET) and TF-target interactions were based on TRRUST and DoRothEA. **a** Enrichment of disease-associated genes for seven autoimmune diseases among the differentially expressed genes by P4. Downregulated DEGs = blue bars, upregulated DEGs = red bars. DEGs were defined comparing cells activated in the presence of P4 as compared to activation alone and represent the total number of uniquely expressed genes at 6 and 24 hrs combined. Dotted line shows p=0.05. The genes that are common between all seven diseases among the differentially downregulated genes by P4 are depicted above the bars. Enrichment p values were calculated by Fisher’s exact test. The TFs with the highest number of interacting genes among the disease-associated differentially downregulated genes for **b** shared genes between the seven different diseases (out of 36 TFs in totalt) and **c** common genes between MS, RA and psoriasis (out of 57 TFs in total). **d-e** Schematic representation of *Stat1* and *Stat3* and their interacting genes. *Stat1* and *Stat3* is significantly downregulated by P4 as are their targets depicted in the figure. DEGs, differentially expressed genes: MS, multiple sclerosis: P4, progesterone: RA, rheumatoid arthritis: TF, transcription factor.

### Disease-associated changes induced by P4 are enriched in targets for T_H_1-associated *STAT1* and T_H_17-associated *STAT3*

To get a more complete understanding of the effect of P4 on disease-associated changes, we sought to identify upstream regulators of the affected disease genes. Using known validated transcription factor (TF)-target interactions from TRRUST^40^ and DoRothEA,^41^ we identified a total of 36 TFs that together regulated 12 out of the 17 shared disease genes. The T_H_1- and T_H_17-associated TFs *STAT1* and *STAT3* were significantly downregulated by P4 and had the highest number of target genes. Furthermore, the targets of STAT1 and STAT3 were significantly enriched among the disease genes (STAT1: n=9, OR: 9.5, p=0.0004; STAT3: n=7, OR: 11.8, p=0.0001) **(Fig. 4b, Supplementary Table 4)**. Next, we focused on MS, RA and psoriasis, that have been shown to markedly improve during pregnancy,^42–44^ in order to more specifically pinpoint which TFs and genes that are affected by P4. There was a total of 62 genes in common between the three diseases that were significantly downregulated by P4 **(Supplementary Table 5)**. A total of 57 different TFs were depicted to regulate 39 of these common genes (2.6 mean number of interactions per TF) where again targets for STAT1 and STAT3 were significantly enriched among the common disease-associated genes (STAT1: n=19, targeting 48% of the common genes, OR: 3.2, p=0.001; STAT3: n=14, targeting 36% of the common genes, OR: 5.6, p=3.0×10^-5^) **(Fig. 5c-e, Supplementary Table 5)**. Thus, the downregulatory effect of P4 on disease-associated genes seems to be primarily mediated through targeting *STAT1* and *STAT3* and their downstream targets.

### Disease-associated transcriptomic changes induced by P4 are mirrored at the proteomics level

To further highlight the role of P4 in the pregnancy-associated improvement of MS, RA and psoriasis, we investigated if the disease-associated transcriptomic changes were also reflected at the protein level. We thereby identified proteins where the known interacting TFs were also significantly affected by P4, and the disease-associated gene as well as the corresponding protein were significantly downregulated by P4. We combined all proteins that were significantly downregulated by P4 at 6, 24 and 72 hrs, to get a global map of the proteomic changes, resulting in a total of 41 DEPs. Focusing on STAT1 and STAT3, we found several disease-associated proteins downstream of STAT1 and STAT3 to be significantly decreased by P4: IL-12β (STAT1), IL-10 (STAT1, STAT3), IL-2 (STAT3), CXCL10 (STAT1), OSM (STAT3) and TFG-β1 (STAT3) **(Table 1)**. Further, we found three additional proteins that were also disease-associated and where their corresponding genes were significantly downregulated by P4: TNF, TNFSF10 and IL-13 **(Table 1)**. Thus, for nine inflammatory-related proteins, we can confirm that the transcriptomic changes induced by P4 also were mirrored at the proteomic levels. Taken together, P4 significantly affects TF expression resulting in downregulation of the interacting disease-associated genes, which in turn dampens the expression of the corresponding proteins.

**Table 1.** Disease-associated genes and proteins and their corresponding transcription factors that are significantly affected by P4.

| Transcription factors | Gene | Protein |
| --- | --- | --- |
| *BACH2, EGR1, FOXP1, FOXP3, NFATC2, RUNX1, STAT3, TOB1* | ↓*IL2* | ↓IL-2 |
| *ATF2, CEBPB, E2F1, EGR1, HMGB2, HSF1, IRF5* | ↓*TNF* | ↓TNF |
| *CEBPB, FOXP3, IRF1, PARP1, STAT1, STAT3, TBX21* | ↓*IL10* | ↓IL-10 |
| *EGR1, HDAC2, IRF1, NFKBIA* | ↓*TNFSF10* | ↓TNFSF10 |
| *EGR1, FOSB, SMAD7, STAT3, USF2* | ↓*TGFB1* | ↓TGF-β1 |
| *IRF1, IRF7, STAT1* | ↓*CXCL10* | ↓CXCL10 |
| *CEBPB, IRF1, STAT1* | ↓*IL12B* | ↓IL-12B |
| *CEBPB, STAT6, TBX21* | ↓*IL13* | ↓IL-13 |
| *STAT3, STAT5B* | ↓*OSM* | ↓OSM |

↓ downregulated by P4 (as compared to activation alone)

## DISCUSSION

In the present study, we demonstrate that P4 significantly dampens T cell activation, thus, providing a compelling explanation for how T cell responses can potentially be regulated during pregnancy when P4 levels are high and immune regulation is required. Transcriptomic analysis revealed that P4 significantly altered the gene expression profile by dampening immune related genes and pathways such as JAK-STAT and T cell receptor signaling. Interestingly, the transcriptomic changes induced by P4 were highly enriched for genes associated with immune-mediated diseases, known to be modulated during pregnancy such as MS, RA, psoriasis and SLE. Further, STAT1 and STAT3 were highlighted as central regulators and several downstream targets were significantly downregulated by P4, also at the protein level, including several well-known and disease-relevant cytokines. Our findings extend previous knowledge about P4 as an immune regulatory hormone and provides further knowledge on how P4 affects CD4^+^ T cell activation and its potential involvement in the pregnancy-associated disease modulation observed for several immune-mediated diseases. Collectively, our findings indicate that P4 plays a significant role in the pregnancy-induced immunomodulation, in particular, by affecting disease-associated T_H_1 and T_H_17 responses. Our findings open up for the exploration of new therapeutic regimes, mimicking the pregnancy situation, in immune-mediated diseases.

Previous studies have investigated the effect of P4 on lymphocyte and CD3^+^ T cell activation.^35, 45–48^ However, the present study is our knowledge not only the first study to directly assess the effect of P4 on human CD4^+^ T cell activation, but also the first study to perform indepth transcriptomic profiling to decipher the effects of P4. Using an *in vitro* system of T cell activation, P4 was found to consistently inhibit T cell activation, as evident from the lowering of the expression of cell surface activation markers and by the profound dampening of immune-related genes and proteins. Further, changes that were specifically related to the actual T cell activation were indeed opposed by P4, *i.e*. genes and proteins upregulated during T cell activation were downregulated by P4 and *vice versa*. T cell activation is indeed a crucial checkpoint for immune regulation in both pregnancy tolerance^49^ and autoimmunity^33, 50^ and administration of P4 has been shown to prevent T cell activation-induced preterm labour and preterm birth in an animal model,^51^ emphasising the importance of P4 in controlling T cell activation during pregnancy.

P4 has been suggested as one potential candidate for the pregnancy-induced immune modulation of several autoimmune diseases, considering its correlation with disease activity during pregnancy. Indeed, pre-treatment with P4 in experimental autoimmune encephalomyelitis (EAE), an animal model of MS, attenuated disease severity and reduced inflammatory responses.^52, 53^ Further, even after EAE onset, P4 administration could significantly reduce disease severity.^54^ Our findings that the P4-induced genes were significantly enriched for disease-associated genes, related to several autoimmune diseases that are known to be modulated during pregnancy, provide additional support for a possible involvement of P4 in this disease modulating activity. Interestingly, there were several genes overlapping between all diseases, indicating potentially shared mechanisms in the pregnancy-induced modulation of these diseases.

It was evident that the predominant effect of P4 on disease-associated genes were primarily related to T_H_1 and T_H_17 responses. Indeed, *STAT1* and *STAT3* are central for promoting T_H_1 and T_H_17 differentiation respectively, both of which have been implicated in the disease pathogenesis of several autoimmune diseases. Loss of STAT3 expression in T cells has for example been shown to confer protection against EAE^55^ and high levels of STAT3 can be used to predict conversion from clinically isolated syndrome to definite MS.^56^ Furthermore, targeting STAT3 has been suggested as a potential therapeutic approach in diseases like MS, psoriasis and RA, where T_H_17 plays a prominent part in the disease processes.^57–59^ P4 has previously been shown to dampen both T_H_1 and T_H_17-related immune responses^20, 60–63^ and is suggested to play a major role in the shift away from T_H_1/T_H_17 in favour of more T_H_2/T_reg_-dominated responses during pregnancy.^19^ However, as far as we know, it has not previously been shown as a possible mechanism for the pregnancy-induced modulation of disease activity. Our transcriptomic findings were validated at the protein level for several highly relevant disease targets downstream of *STAT1* and *STAT3*. Targeting IL-12β (IL12p40), which constitutes a crossroad between T_H_1-associated IL-12 and T_H_17-associated IL-23, is successfully implemented as treatment for psoriasis and is undergoing clinical trials for several other immune-mediated inflammatory diseases.^64^ Clearly, IL-12β plays a profound role in disease and the dampening effect of P4 on this gene and subsequent protein expression could thus constitute a possible avenue in the modulation of disease by P4. It is noteworthy that P4 did not only dampen pro-inflammatory but also anti-inflammatory genes and proteins such as IL-10 and TGF-β, indicating that P4 induces a rather general downregulation of immune responses. It is well-known that anti-inflammatory responses are often induced as a negative feedback mechanism during inflammation to help prevent excessive inflammatory responses and help restore tissue homeostasis. Speculatively, as P4 dampens immune responses, it is only natural to assume that lesser anti-inflammatory responses would follow as a result. This is exemplified through the intricate relationship between STAT3 and IL-10, where the *IL10* promoter contains a specific binding motif for STAT3, which appears to control the expression of IL10.^65^ Indeed, increased expression of STAT3 in T cells from patients with SLE promoted IL-10 expression^66^ and similar associations have been proposed between STAT3 and TGF-β as well.^67, 68^ Taken together, the profound dampening of activation in CD4^+^ T cells, including several T_H_1 and T_H_17-associated pathways, indicate that P4 is able to inhibit proinflammatory responses.

In light of our findings, it is tempting to speculate of a potential beneficial therapeutic effect of P4 in autoimmune diseases. In humans, although circumstantial, worsening of symptoms prior to menstruation, when P4 levels are lower, have been reported in both RA and MS.^69–71^ Despite that a previous clinical trial using high dose progestins (synthetic progestogens) failed to prevent post-partum relapses in patients with MS,^72^ it does not exclude the possibility of using progestogens as potential treatment. Since the period after delivery is associated with overall dramatic changes, not only in hormone levels but also in the reversal of the immune system to the pre-pregnancy state, it is not so surprising that no clear effects were observed. Progestogens still represent a very attractive treatment option since they can vary in their androgen, progesterone and glucocorticoid receptor affinity and they could therefore, theoretically, have very diverse effects on different aspects of the disease pathogenesis. Thus, it could be rewarding to look further into the therapeutic potential of progestogens.

One potential limitation of our study is that we investigated the effect of P4 in healthy females. To the best of our knowledge, there are no studies that have specifically evaluated if patients with autoimmune diseases would respond differently to P4 than healthy individuals. However, when considering P4 as a potential treatment option this needs to be further investigated. Further, the responsiveness to P4 could vary as the P4-related receptor expression could vary throughout the menstrual cycle.^73, 74^ The women included in this study were sampled evenly across the menstrual cycle which should minimise the bias in differences in receptor expression. Furthermore, all included women responded to P4 in a consistent way. Another remaining caveat which we have yet to address is understanding the precise mechanisms by which P4 operates. This is complicated by the fact that P4 is a promiscuous hormone that can bind several targets. There are conflicting results regarding the expression of different P4-related receptors in CD4^+^T cells.^23, 35, 48, 75^ Further, whether P4 works mainly by slower genomic or more rapid non-genomic actions through nuclear or membrane receptors respectively, remains to be settled. Interestingly, lymphocytes from pregnant as compared to non-pregnant women appear to have increased sensitivity to P4,^76^ which has been suggested to be attributed to a more active immune system during pregnancy triggered by the presence of the semi-allogenic foetus. Furthermore, P4 has also been shown to interact with the glucocorticoid receptor (GR)^77, 78^ and the synthetic corticosteroid dexamethasone has been shown to exert similar effects on T cell differentiation as P4.^63^ Considering the concentrations used here, one cannot rule out that some of the observed effects are derived from binding to the GR.

In summary, we showed that the presence of P4 significantly reduced activation of CD4+ T cells and induced large transcriptomic changes in the activated cells. Most prominently P4 downregulated immune responses associated with the T cell activation and the genes affected by P4 were significantly enriched for disease-associated genes of immune-mediate diseases that are known to be modulated during pregnancy. The most pronounced effect of P4 on disease-associated genes was primarily related to T_H_1 and T_H_17 responses. We conclude that our study supports a role for P4 in the immune-modulation induced during pregnancy and that P4 should be further evaluated as a potential treatment option in T-cell mediated disease.

## MATERIALS AND METHODS

### Study subjects

Blood samples were collected from thirteen healthy female volunteers (median age 32, 25-43 yrs), recruited among students and personnel at Linköping University and Linköping University Hospital, Sweden. Informed consent was obtained prior to sample collection and the study was approved by the Regional Ethics Review Board in Linköping (Regionala etikprövningsnämnden i Linköping), Sweden, (approval number: M39-08). None of the women were using hormonal contraceptives or taking any other medications at the time of inclusion. The time points of sample collection were evenly distributed across the menstrual cycle (assuming a 28-day menstrual cycle; seven women were in the luteal and six in the follicular phase).

### Isolation of CD4^+^ T cells

Peripheral blood mononuclear cells (PBMCs) were isolated by gradient centrifugation using Lymphoprep™ (Axis-Shield, Oslo, Norway) and washed thrice in Hank’s Balanced Salt Solution (Life Technologies, Darmstadt, Germany). Magnetic activated cell sorting (MACS) was used to isolate CD4^+^ T cells. The PBMCs were resuspended in MACS buffer (phosphate buffered saline, PBS; Medicago, Uppsala, Sweden) supplemented with 2mM EDTA (Sigma Aldrich, Saint Louis, MO, USA) and 0.5% fetal bovine serum (FBS; Thermo Fisher Scientific, Waltham, MA, USA) and the CD4^+^ cells were isolated by positive immunomagnetic selection using MS columns and a mini MACS separator (Miltenyi Biotec, Bergish Gladbach, Germany) according to the instructions provided by the manufacturer. The purity of the isolated CD4^+^ T cells was assessed by flow cytometry (median purity 98.5%, range 97.6-99.0%).

### Pre-incubation with progesterone

The isolated CD4^+^ T cells were pre-incubated with 10, 30 and 50 μM of P4 (water-soluble; Sigma Aldrich) or without (cell culture media alone). The cells were plated in 24-well flat-bottom plates (Costar™; Corning Inc, Corning, NY, USA) at 1.0×10^6^cells/ml, in a 1 ml final volume/well of Isocove’s Modified Dulbecco’s Medium (IMDM; Invitrogen, Carlsbad, CA, USA) supplemented with L-glutamine (292 mg/l; Sigma-Aldrich), MEM non-essential amino acids 100X (10 ml/l; Gibco^®^), penicillin (50 IE/ml), streptomycin (50 μg/ml; Cambrex-Lonza, Basel, Switzerland), and sodium bicarbonate (3.024 g/l; Sigma-Aldrich) and 5% FBS at 37°C and 5% CO_2_. After 20 hrs, the cells were removed from the plates, centrifuged and resuspended in cell culture media prior to *in vitro* stimulation. See **Fig. 1a** for an overview of the experimental design of the study.

### *In vitro* activation of CD4^+^ T cells in the absence or presence of P4

Twenty-four-well flat bottom plates (Costar™) were coated with 0.1 μg/ml of low endotoxin anti-CD3 and anti-CD28 antibodies (clone UCHT1, clone YTH913.12; Bio-Rad AbD Serotec Limited, Hercules, CA, USA) or with PBS alone for 20 hrs at 4°C followed by washing thrice in PBS. The concentration of antibodies was chosen based on titration experiments where 0.1 μg/ml resulted in moderately increased cell surface expression of the early T cell activation marker CD69. The CD4^+^ T cells pre-incubated without P4 were cultured unactivated or activated (with anti-CD3/CD28 antibodies), whereas the CD4^+^ T cells pre-incubated with P4 were activated in the presence of the same concentrations as during the pre-incubation for 6-24-72 hrs and subsequently processed for flow cytometry, RNA extraction or measuring of secreted proteins. Cells cultured for 72 hrs were only used for protein measurements. Briefly, after culturing, the supernatants were collected and a portion the cells were used for flow cytometry analysis and the rest lysed and homogenized in buffer RLT Plus (Qiagen; Hilden, Germany) supplemented with 143 mM β-mercaptoethanol (Sigma Aldrich) and homogenized using syringe and needle according to the instructions provided by the manufacturer. The lysates were stored at −70°C before extraction.

### Evaluation of activation status with flow cytometry

The CD4^+^T cells were resuspended in LIVE/DEAD™ Fixable Dead Cell Stain (Invitrogen), diluted 1:500 in PBS+0.1%FBS and stained with mouse anti-human CD4-FITC (clone SK3), CD69-APCCy7 (clone FN50), CD25-PE (clone 2A3) and CD3-APC (clone SK7; all from BD Biosciences, Franklin Lakes, NJ, USA). The cells were incubated in the dark for 15 minutes at room temperature and washed in PBS+0.1%FBS prior to flow cytometry analysis. 10 000 CD4^+^ T cells were collected and analyzed using FACS Canto II (BD Biosciences) and Kaluza software version 2.1 (Beckman Coulter, Brea, CA, USA). The cells were gated according to forward (FSC) and side scatter (SSC) and further defined as CD3^+^CD4^+^ **(Supplementary Fig. 2)**. The cut-off value for CD69 expression was based on its expression in unactivated CD4^+^T cells and CD25 expression was set based upon the contour of the identified negative and positive populations. Fold change of CD69 and CD25 expression was calculated based on expression of the activation markers on the cells activated without P4 (expression activated cells with P4/expression activated cells without P4). All data were analyzed using GraphPad Prism version 8.0.1 (San Diego, CA, USA). The majority of the data was normally distributed and therefore analyzed using one-way ANOVA with Dunnett’s multiple comparison test. Data are expressed as mean and standard deviation and p-values ≤ 0.05 were considered statistically significant.

### RNA sequencing

RNA was extracted using the AllPrep DNA/RNA Mini kit (Qiagen) according the protocol provided by the manufacturer and RNA was eluted with 30 μl of RNase-free water. The concentration of RNA was determined using Nanodrop^®^ ND-1000 spectrophotometer (Nanodrop Technologies Inc; Wilmington, DE, USA). The RNA quality was controlled using Agilent RNA 6000 Nano Kit (Agilent Technologies, Santa Clara, CA, USA) on an Agilent 2100 Bioanalyzer instrument (Agilent Technologies). RNA integrity numbers (RIN) were 9.5±0.4 (mean±SD). Libraries were constructed with TruSeq Stranded mRNA (Illumina, San Diego, CA, USA), which have been adapted to run on an Agilent BRAVO robot (Agilent Technologies) with 440 ng of mRNA as starting material per sample. Briefly, poly-A containing mRNA was isolated using poly dT-coated beads and broken down into 150-400 base pair fragments by chemical fragmentation and converted to cDNA with reverse transcriptase and random primer. Enzymes and shorter fragments were removed using AMPure XP beads (Beckman Coulter, Indianapolis, IN, US). The remaining fragments were adenylated followed by ligation of index-adapters and amplified with PCR. Samples were barcoded, pooled and sequenced on the Illumina NovaSeq 6000 platform with a S1 flowcell and sequenced PE2×101 bp. An average of 48.9 million reads were obtained per sample (48.9×10^6^±10.5×10^6^, mean ± SD). Per-cycle base call (BCL) files were demultiplexed and converted to FASTQ using bcl2fastq v2.19.1.403 from the CASAVA software suite (Illumina). Library preparation and sequencing was carried out at the National Genomics Infrastructure, Science for Life Laboratories, Stockholm. Paired samples from a total of eight individuals (n=38 samples in total) were used for RNA-seq.

### RNA-seq data analysis

The FASTQ files were processed with TrimGalore! to remove adapter contamination and trimming of low-quality regions. Paired-end reads were aligned and mapped to the Ensemble human reference genome GRCh37 (Genome Reference Consortium Human Build 37) using STAR (version 2.5.3.a).^79^ Gene read counts were generated using StringTie (version 1.3.3).^80^ Data was processed in R Studio (Version 1.1.456; Boston, MA, USA) using the edgeR^81, 82^ and limma packages.^83, 84^ The mapped reads were filtered for lowly expressed genes (genes with counts per million (cpm) >1 in at least 3 replicates were kept) and used for further analysis. The filtered gene counts were normalized by using the trimmed mean of M-values (TMM) via calcNormFactors in edgeR. Voom-transformation was applied prior to differential expression analysis. An FDR (Benjamini-Hochberg) of 0.05 was used as a threshold for DEGs. Differential expression induced by P4 was calculated comparing CD4^+^T cells activated in the presence or absence of P4. A summary of the RNA-seq data analysis can be seen in **Supplementary Fig. 3**.

### Pathway, gene set enrichment analysis and gene overlap

To explore the biological relevance of the DEGs, a gene set enrichment (GSE) analysis using *gseKEGG* and pathway analysis using *enrichKEGG* from Clusterprofiler^85^ based on the Kyoto Encyclopedia of Genes and Genomes (KEGG)^86, 87^ was performed. For GSE analysis, a pre-ranked list of P4-induced DEGs (based on logFoldChange (logFC), n=3339 genes at 6 hrs and n=3725 genes at 24 hrs) was used as input, where genes that lacked Entrez gene ID were removed prior to analysis. An enrichment score (ES) was calculated for all gene sets and normalized for gene set size (normalized enrichment score; NES) over a mean distribution of 1000 permutations. The minimum gene set size was set to 20 and the maximum to 200 genes, thereby excluding small and very large generic data sets. An adjusted p-value ≤ 0.05 (Benjamini-Hochberg) was considered statistically significant. Gene overlap between different gene sets was performed using the GeneOverlap package in R (version 4.0).^88^

Measurement of secreted proteins in culture supernatants using Proximity Extension Assay In order to verify central transcriptomic changes on the protein level, culture supernatants collected after 6-24-72 hrs were analysed for 92 inflammation-associated proteins (Olink inflammation panel; https://www.olink.com/products/inflammation/) with multiplex PEA at the Clinical Biomarkers facility, Science for Life Laboratory, Uppsala University, SE-751 85 Uppsala. Briefly, 1 μl of cell supernatant was incubated with matched pairs of antibodies linked to unique oligonucleotide (proximity probes) specific for each biomarker to be measured. Close proximity of the probes bound to their targets results in hybridization which, with the addition of DNA polymerase, extends the oligonucleotides, creating a DNA amplicon that can be detected and quantified by quantitative real-time PCR.^37^ Four internal controls were included for quality control and for data normalization. Data was expressed as normalized protein expression (NPX), an arbitrary unit in log_2_ scale. Values below the detection limit were assigned half the value of the lowest detection limit. Proteins that were detected in a least 50% of the samples at each time point were included in the statistical analysis with the following exceptions: CCL20 (40% detectable at 24 hrs), TGF-β1 (40% detectable at 72 hrs) and LIF (47% at 24 hrs) where more than half of the samples in the activation alone group was detectable (8-10 out of 10) and significantly higher than the unactivated samples **(Supplemental Table 2)**. Statistical differences were determined using Friedman test and Banjamini-Hochberg to correct for multiple comparisons. An adjusted p-value of ≤0.05 was considered as statistically significant. All data was analysed with GraphPad Prism version 8.0.1.

### Enrichment analysis of disease-associated genes

Disease-associated genes for Hashimoto’s (n=141), Graves’ disease (n=245), MS (n=801), psoriasis (n=580), RA (n=1273), SLE (n=817) and SS (n=) were derived from DisGeNET.^38^ All disease genes that were not present in the background (all genes detected by RNAseq, n=14363) were discarded from downstream analysis. Enrichment analysis of disease-associated genes among the genes affected by P4 were performed using GeneOverlap in R that uses Fischer’s exact test to compute p values and odds ratio for the overlaps.^88^ The P4-induced DEGs that were used comprised the uniquely expressed genes combining the DEPs from both 6 and 24 hrs (in total n= 2563 downregulated and n= 2403 upregulated genes).

### TF-target interactions

TF-target interactions were derived using TRRUST^40^ and DoRothEA^41^. The total number of interactions from TTRUST was 9396 and from DoRothEA (including interactions with confidence score A and B) was 6620 but limiting the interactions to TFs that were significantly affected by P4 resulted in a total number of 3124 interactions, combining both TTRUST and DoRothEA. First we identified TFs that were significantly affected by P4 and then only included those TF-target interactions where the target was also significantly downregulated by P4 and disease-associated as established by the common genes between all diseases or between MS, psoriasis and RA. Enrichment of targets for *STAT1* and *STAT3* was computed using Fisher’s exact test where targets among the disease-associated genes that were downregulated by P4 was compared to the targets of the corresponding TFs among all genes downregulated by P4. For TF-mRNA-protein analysis, we combined the DEPs derived from the measurement of the expression of 92 inflammation-related proteins in culture supernatants at 6-24-72 hrs (n=41). These proteins were mapped to the corresponding mRNA and, in extension, to the known interacting TFs.

## DATA AVAILABILITY

Data will be deposited on GEO upon publication.

## ACKNOWLEDGEMENTS

The authors acknowledge the National Genomics Infrastructure (NGI) in Stockholm funded by Science for Life Laboratory, the Knut and Alice Wallenberg Foundation and the Swedish Research Council, and SNIC/Uppsala Multidisciplinary Centre for Advanced Computational Science for assistance with massively parallel sequencing and access to the UPPMAX computational infrastructure. The authors would also like to acknowledge the support of the Clinical biomarker facility at SciLifeLab Sweden for providing assistance in protein analyses.

## AUTHOR CONTRIBUTIONS

SH, JR and GP collected samples and performed cell culture experiments and flow cytometry. SH, OR and RM performed pre-processing, analysis of the RNA-sequencing data and bioinformatics analysis. JE, MG and MCJ contributed to the study design and overall supervision of the study. SH was responsible for preparation of figures and the writing of the manuscript with the support of MG, JE and MCJ. All authors read and approved the final manuscript.

