## Supplementary Figures for "Progesterone specifically dampens disease-associated T_H_1- and T_H_17-related immune responses during T cell activation *in vitro*"

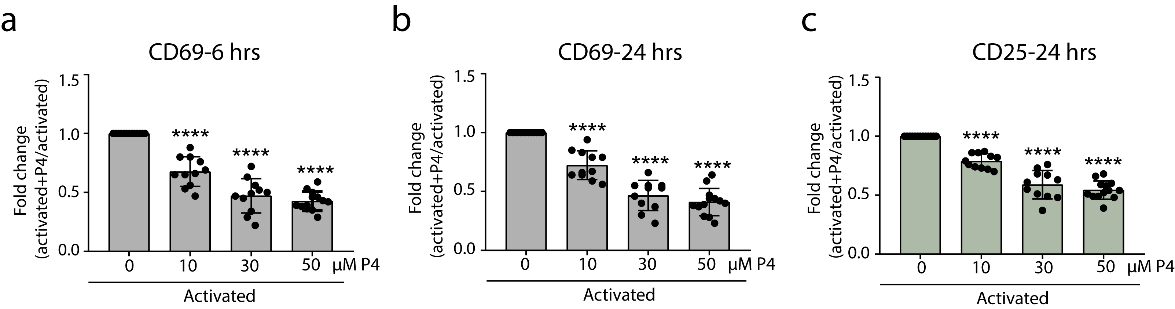
**Fig. S1.** Fold change in activation markers in CD4^+^ T cells activated in the presence of P4. Isolated CD4^+^ T cells were activated in the presence or absence of 50 µM of P4 for 6 and 24 hrs. Activation status was evaluated by flow cytometry using the surface activation markers CD69 (6 and 24 hrs) and CD25 (24 hrs). Figure shows mean ± standard deviations of the fold-change in expression of the activation markers comparing activation in the presence of P4 as compared to activation alone. Statistical differences were determined using one-way ANOVA with Dunnett’s multiple comparison test. n=13, ****p≤0.0001. P4: progesterone.

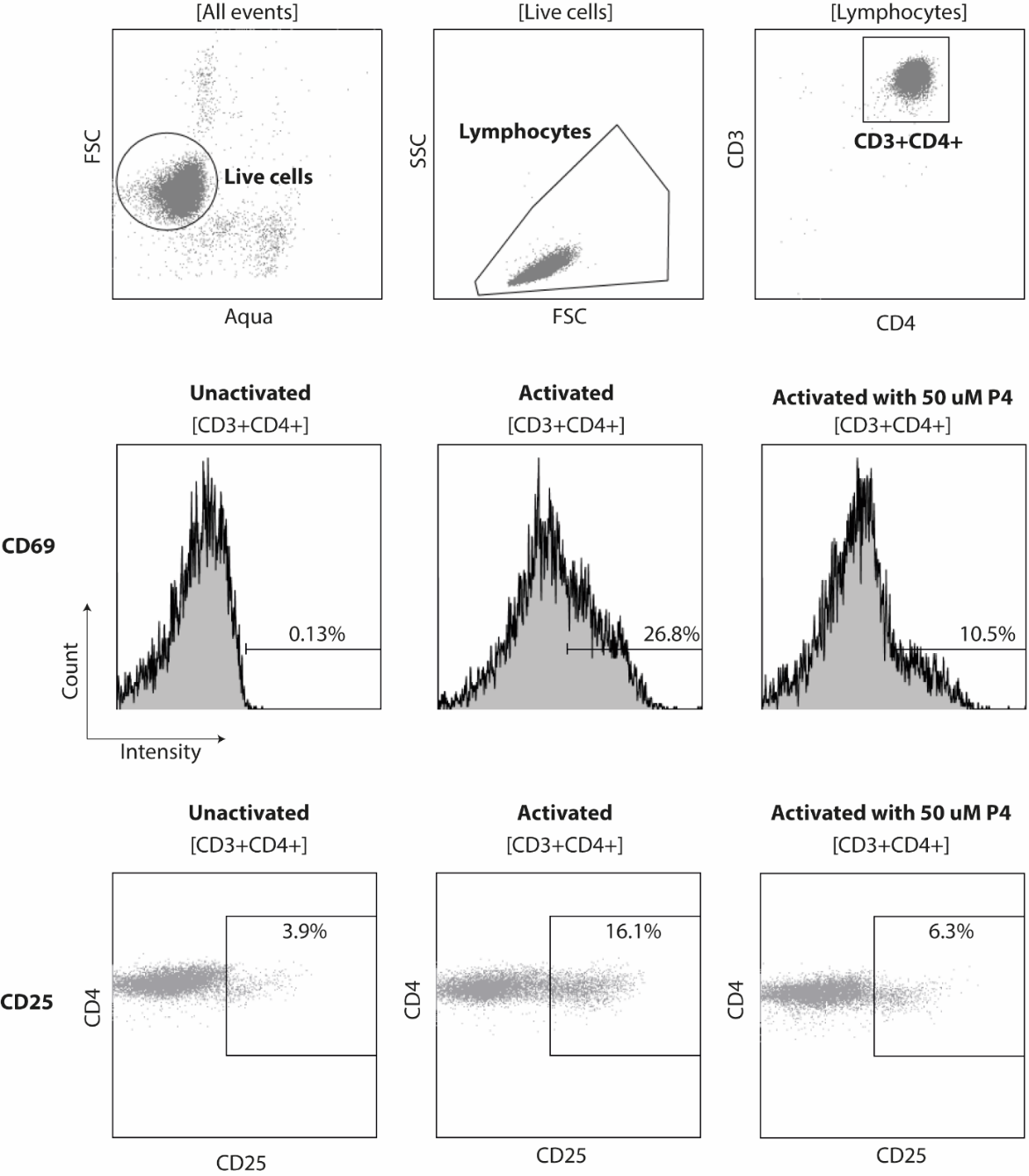
**Fig. S2.** Gating strategy for analysis of T cell activation markers. Cells were gated as live cells based on negative expression of Aqua Live/Dead stain and gated based on forward and side scatter. CD3^+^CD4^+^ T cells were analysed for the expression of CD69 (6 and 24 hrs) and CD25 (24 hrs) and the percentage of cells expression the markers after activation was set based on the expression in the unactivated cells. Figure shows one representative sample. FSC: forward scatter, P4: progesterone, SSC: side scatter.
